# A molecular rack and pinion actuates a cell-surface adhesin and enables bacterial gliding motility

**DOI:** 10.1101/267583

**Authors:** Abhishek Shrivastava, Howard C. Berg

## Abstract

The mechanism for bacterial gliding is not understood. The gliding bacterium *Flavobacterium johnsoniae* is known to have an adhesin, SprB, that moves along the cell surface on a spiral track. When cells are sheared by passage of a suspension through thin tubing, they stop gliding but can be tethered by addition of an anti-SprB antibody. Tethered cells spin about 3 Hz. We labeled the Type 9 secretion system (T9SS) with a yellow-fluorescent-protein (YFP) fusion of GldL. When labeled cells were tethered, a yellow fluorescent spot was found near the rotation axis, which shows that the motor that drives the rotation localizes with the T9SS. The spiral track was labeled by following the motion of Cy3 attached to SprB via an antibody. The distance between the rotation axis and the track was determined by a measurement involving both labels, YFP and Cy3, yielding 90 nm. If a rotary motor spins a pinion of radius 90 nm 3 Hz, a spot on its periphery will move 1.5 μm/s, the speed at which cells glide. We suggest that the pinion drives a flexible tread that carries SprB along a track fixed to the cell surface. Cells glide when such an adhesin adheres to the solid substratum.

## Introduction

Rod-shaped gliding bacteria move in a manner similar to a self-propelled screw, i.e., they roll along their long axis as they move forward on an external surface^1,2^. Bacteria related to *Flavobacterium johnsoniae*, which is the fastest of all known gliders, are present in diverse environments such as the human oral microbiome^3^, fish scales^4^, water bodies^5^, and plant rhizosphere^6^ where they glide over and colonize their preferred surfaces. How the molecular players form the machinery that actuates this motion is not clearly understood. *Flavobacterium johnsoniae* is now the organism of choice for studies of this process, because rates of movement are high and much of the genetics is known^7,8^. A mobile cell-surface adhesin, SprB, has been identified that plays a central role in gliding^9,10^. Tracking of SprB in 3D space has revealed the presence of a spiral cell-surface track on which SprB moves^1^. Cells subjected to viscous shear stop gliding, but they can be tethered to an external surface using anti-SprB antibody. Tethered cells pinwheel around a fixed axis, suggesting that a rotary motor that generates high torque is a part of the gliding machinery^11^.

Bacterial motility machines have external components, so are coupled with protein secretion systems. The flagellar motor is associated with the Type 3 secretion system^12,13^, and the Type 4 pili motor is associated with the type 2 secretion system^14-16^. In *Flavobacterium*, the Type 9 secretion system (T9SS) is required for the secretion of SprB, and cells lacking the T9SS are non-motile^17^. T9SS spans the inner and outer membranes and the periplasmic region. The T9SS has an outer membrane barrel made up of a protein SprA. It is one the largest transmembrane β barrels known in biology^18^. Structural data suggest that in the periplasmic region, the T9SS forms at least one ring-shaped structure that contains GldK, while GldM spans the periplasmic region^19, 20^. GldL is a cytoplasmic membrane protein and is one of the core T9SS proteins^17^. Figures that summarize the current proposed macromolecular arrangement of T9SS can be found in both refs. 18 and 19. Our results suggest that T9SS is associated with the rotary component of the gliding motor and drives a tread carrying the adhesin SprB along a track fastened to the rigid framework of the cell wall.

## Results

### The axis of rotation is localized near GldL

GldK, GldL, GldM, GldN, SprA, SprE, SprF and SprT are the core T9SS proteins required for the transport of SprB to the external surface. Δ*gldL* mutants do not have SprB on the cell surface. They are not able to stick to a glass surface and do not display motility^17^. We found that tethered cells of Δ*gldL* mutants complemented by GldL-YFP were able to bind to glass and exhibited wild-type levels of cell rotation (Movie 1). This suggests that the rotary motor is fully functional in this strain.

Fluorescent GldL-YFP appeared as spots that were fixed in the frame of reference of a cell (Fig. 1A & Movie 2). Using ImageJ and custom MATLAB codes C1-C3 that fit a Gaussian to each GldL spot, the positions of GldL foci from 42 cells were determined. Custom MATLAB codes used in this article are freely available on GitHub https://github.com/Abhishek935/Molecular-Rack-and-Pinion. The GldL spots appeared to be localized at random within a cell, with the average distance between two neighboring spots about 1.5 µm (Fig. S1). The peak of a Gaussian fit to a frequency distribution of number of foci per cell as measured from 42 cells is 3.47 with a S.D. of 0.87 (Fig. 1B). The Total Internal Reflection Fluorescence (TIRF) microscope used for these measurements strongly illuminates only the cylindrical face of the cell adjacent to the glass with a penetration depth around 100 nm. A similar distribution of GldL is expected on the opposite cylindrical face. Hence, the total number of GldL foci per cell is about 7.

**Figure1.**
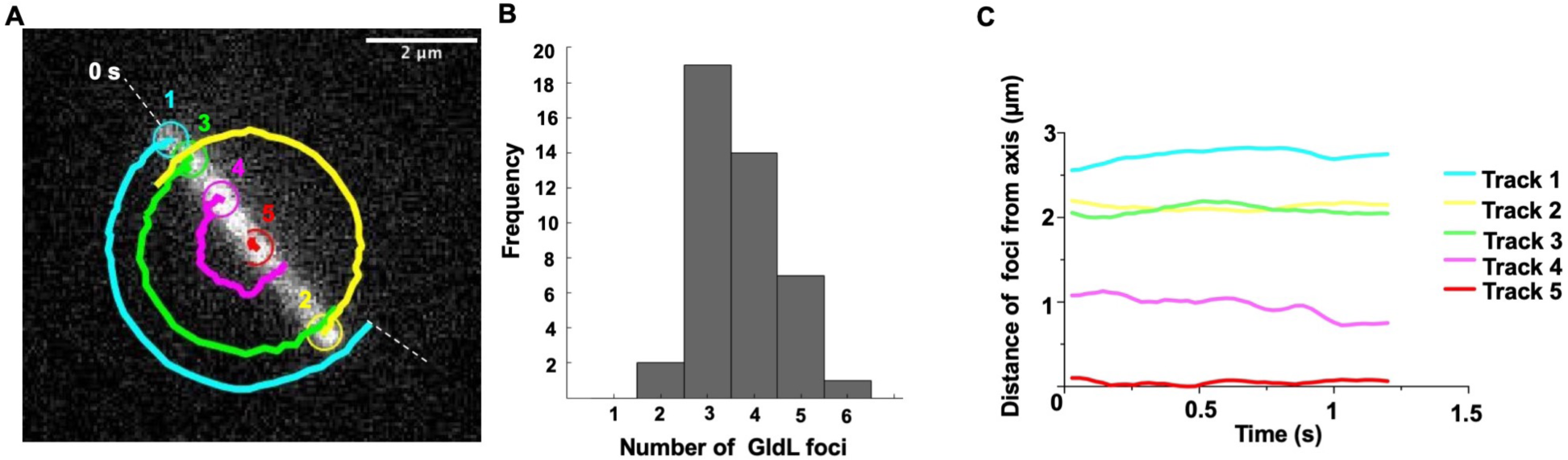
The axis of rotation is found near a GldL spot. **(A)** One tethered cell with 5 GldL foci is shown, with one foci numbered 5 close to the axis of rotation; we call it the ‘nearest neighbor’ GldL. The other foci execute circular arcs as the cell rotates. The foci were spaced about 1.4 µm apart. **(B)** The distribution of numbers of GldL foci per cell for 42 cells. About 4 GldL foci appeared per cylindrical face of a cell. (**C**) Distances of the GldL foci of the tethered cell shown in Fig.1A from the axis of rotation plotted as a function of time.

Using tethered cells of Δ*gldL* background complemented with GldL-YFP, the position of the axis of rotation was determined by adding successive images of an image stack of a rotating cell and fitting the new matrix that contains the sum of images with a Gaussian that has the X-Y coordinates as position and the Z coordinate as intensity. Code C4 was used to add successive images. Given that the axis of rotation remains fixed over all image frames of a rotating cell, for a matrix that contains the sum of stacks, the intensity is highest at the axis of rotation. All GldL foci in a rotating cell were tracked over several frames. All but one GldL spot per tethered cell showed a circular trajectory (Fig. 1A). The foci that did not move with an apparent circular trajectory were within a distance range of few hundred nanometers from the axis of rotation (Fig. 1C). This spot was named the ‘nearest neighbor’ GldL.

Similar measurements were made for each image frame from image stacks of 8 rotating tethered cells with 60 frames on average per image stack. This provided a total of 484 measurements. The standard deviations for the Gaussian fits to the images of the rotation axes and the nearest-neighbor GldL foci bloom up to resolving power of the microscope. The position of nearest neighbor GldL relative to the axis of rotation is shown for all 8 cells in Fig. 2A, with the axis of rotation normalized to be at the origin. The position of nearest neighbor GldL relative to the axis of rotation, one cell at a time, is shown in Fig. S2. The peak of a Gaussian fit to a frequency distribution of distance between the nearest neighbor GldL and the axis of rotation for all 484 measurements was 109.7 nm with a S.D. of 68.5 nm (Fig. 2B). These measurements tell us that the axis of rotation was found near a GldL spot, as expected if GldL is part of the rotary motor. However, we do not know whether GldL is centered on the rotation axis.

**Figure2.**
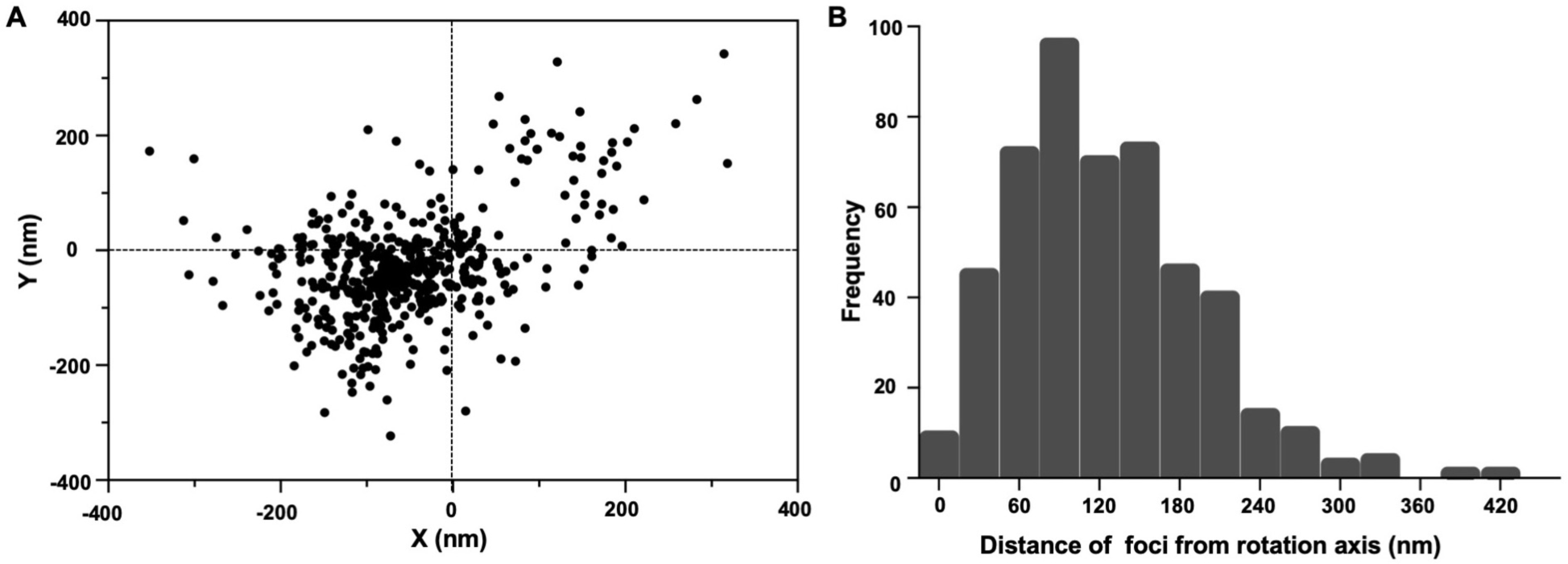
Analysis of the position of GldL spots. **(A)** A scatter plot of the positions of the nearest neighbor GldL relative to the axis of rotation of 8 tethered cells, with 60 frames on average imaged for each cell, resulting in 484 measurements. For each cell, the axis of rotation was normalized to be at the origin. **(B)** Frequency distribution of the distance of the nearest neighbor GldL from the axis of rotation of 8 tethered cells measured for a total of 484 image frames with about 60 image frames per cell. The peak of a Gaussian fit to this distribution is 109.7 nm with a S.D. of 68.47 nm.

### Cell rotation and SprB motion utilize the same power source

A preparation of tethered cells was exposed to 10 µM carbonyl cyanide-m-chlorophenylhydrazone (CCCP), which is known to abolish the protonmotive force that acts across the inner cell membrane^21^. The cells stopped rotating after the addition of CCCP and started rotating again after its removal (Fig. S3, S4 and Movie 3). The loss of rotation was rapid and the recovery was gradual, taking about 4 times longer, as expected if time was required to wash CCCP out of the cells. These results are consistent with previous observations where motion of SprB was stopped after the addition of CCCP and was restored after washing it away^10^. This suggests that both cell rotation and SprB translocation utilize the same fuel.

*MreB, a protein that helps to build the cell wall, is involved in the gliding motility of a rod-shaped bacterium Myxococcus xanthus*^*22*^. A22, a drug that alters the motion of MreB and stops the gliding of *M. xanthus*^*22*^, did not affect the gliding of *F. johnsoniae* (Fig. S5). As in *F. johnsoniae*, binding of a cell-surface adhesin to an external surface results in gliding of *M. xanthus*. However, there are obvious differences between the gliding of the two types of bacteria. *M. xanthus* uses different motility proteins and glides about 60 times more slowly than *F. johnsoniae*.

### The track on which SprB moves is close to the gliding motor

While the motor labeled by GldL-YFP is fixed on the cell surface, the SprB adhesin moves along the length of the cell. How rotation might be coupled to linear motion is a puzzle. The Δ*gldL* strain is complemented for rotary motor function by the addition of GldL-YFP (Movie 1). However, long distance motion of the cell and the adhesin SprB is not fully restored i.e. some cells and beads attached to SprB move very short distances (Movie 1). This suggests that the C-terminal region of GldL might play a role in coupling the rotary motor with the motion of SprB. In order to study long distance movement of SprB in a *gldL-yfp* strain, a *gldL-yfp*-containing plasmid was added to wild-type cells to form a hybrid motor containing both GldL and GldL-YFP subunits. Western blot analysis suggests that GldL expressed from the chromosome and pCP23 is of similar cellular concentration^17^. A Cy3 tag was attached to SprB via an antibody and the positions of both SprB and GldL were determined within same cells (Fig. 3A, S6, and Movie 4). The shortest distance between the trajectory of SprB labeled with Cy3 and motors labeled with GldL-YFP was measured for 42 motors from 10 cells. The average positions of SprB labeled with Cy3 and GldL-YFP (motor) were determined by fitting Gaussians to respective foci from all frames of an image stack. The positions of motors were at a distance (*d*) of 90.9 nm with a S.D. of 63.7 nm from the SprB trajectory (Fig. 3B).

**Figure 3.**
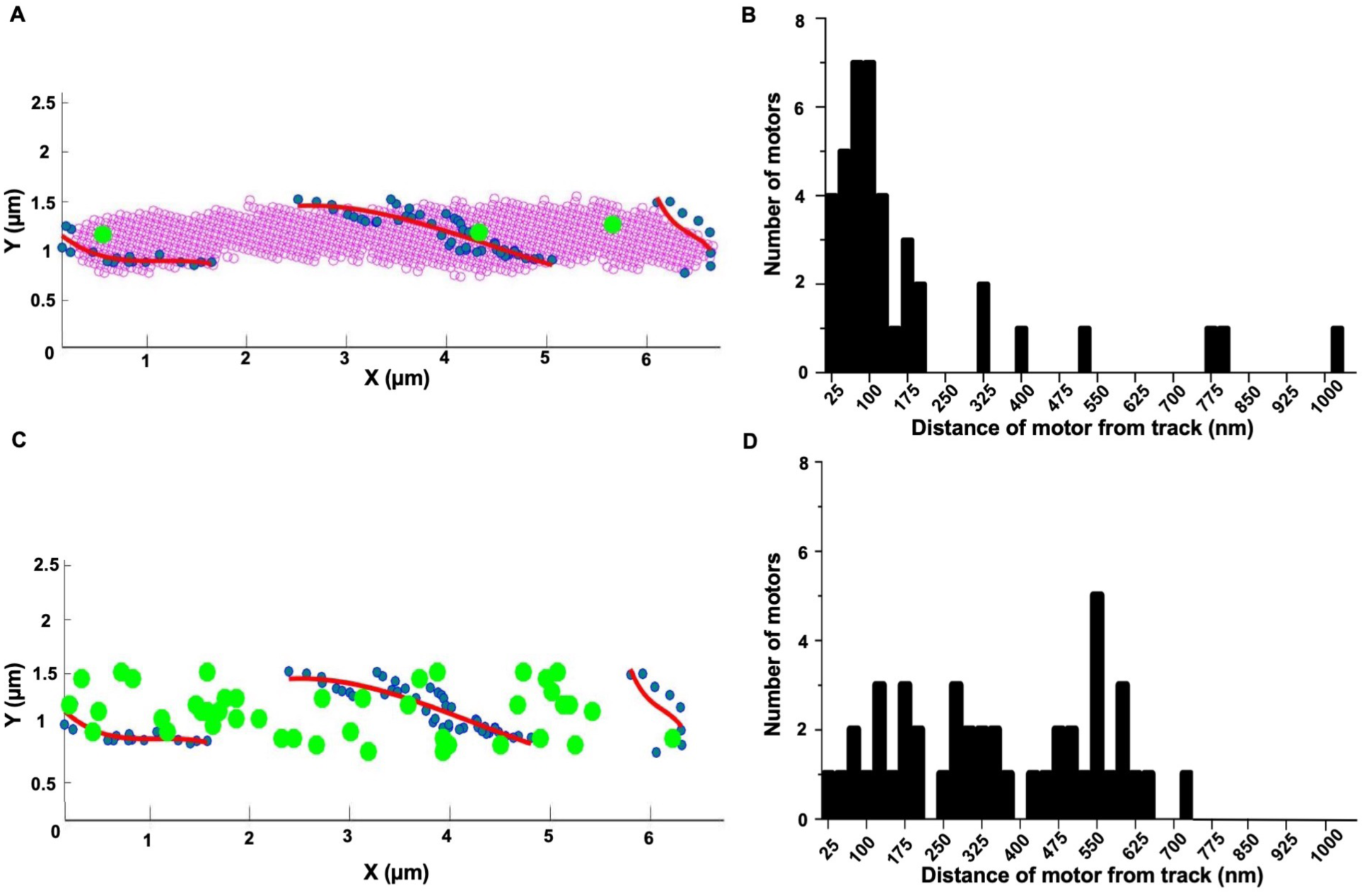
GldL localizes near the track on which SprB moves. **(A)** Co-localization of GldL (green circles) with the trajectory of SprB (blue dots and red line). GldL remains fixed relative to the cell. SprB moves along the trajectory shown in red. Blue dots mark the positions of SprB determined at different time points. Red lines are second-order polynomial fits to the blue dots. GldL was labeled with YFP and SprB with Cy3. The position of SprB and GldL-YFP from the same cell is determined from Movie 4 and Fig. S6 (B) respectively. The images were rotated and translated to determine the positions of SprB and GldL-YFP relative to the cell. **(B)** Frequency distribution of the distance of 42 motors from SprB trajectories for 10 cells, showing a peak at a distance of 90.9 nm with a S.D. of 63.7 nm. **(C)** A simulation in which 42 motors were localized at random on one cell, with the positions of the motors (green) and the track (red). **(D)** A frequency distribution of distances between the motors and the track.

As a control, a simulation in which 42 motors were randomly localized in one cell was performed, together with the SprB trajectory from one cell. This simulation was carried out using a random number generator in MATLAB and similar boundaries as that of a cell in an image plane were used. The shortest distances between the motors and the track were measured for the simulation. The distribution for these distances was broad and did not show a discernible peak near 90 nm (Fig. 3C and 3D).

Tracks carrying SprB did not loop around GldL spots. This argues against a model in which chains carrying SprB are driven by sprockets^7^. A viable alternative, explained below, involves racks and pinions.

### A model for gliding is suggested in which a pinion attached to the rotary motor’s drive shaft engages a rack (a tread) that slides along a track fixed to the rigid framework of the cell wall

A cartoon illustrating this idea is shown in Fig. 4A. A pinion (a circular gear) and a rack (a straight element with teeth matching those on the gear) is a common device for converting rotary motion into linear motion. If the motor rotates at frequency *f* and the radius of the pinion is *r*, then the rack will move at velocity *v* = 2*πrf*, which is the gliding speed, about 1.5 µm/s. Given the maximum *f* that the rotary motor exhibits of about 3 Hz^7^, we find *r* = *v*/2*πf* = 80 nm, which is on the mark.

**Figure 4.**
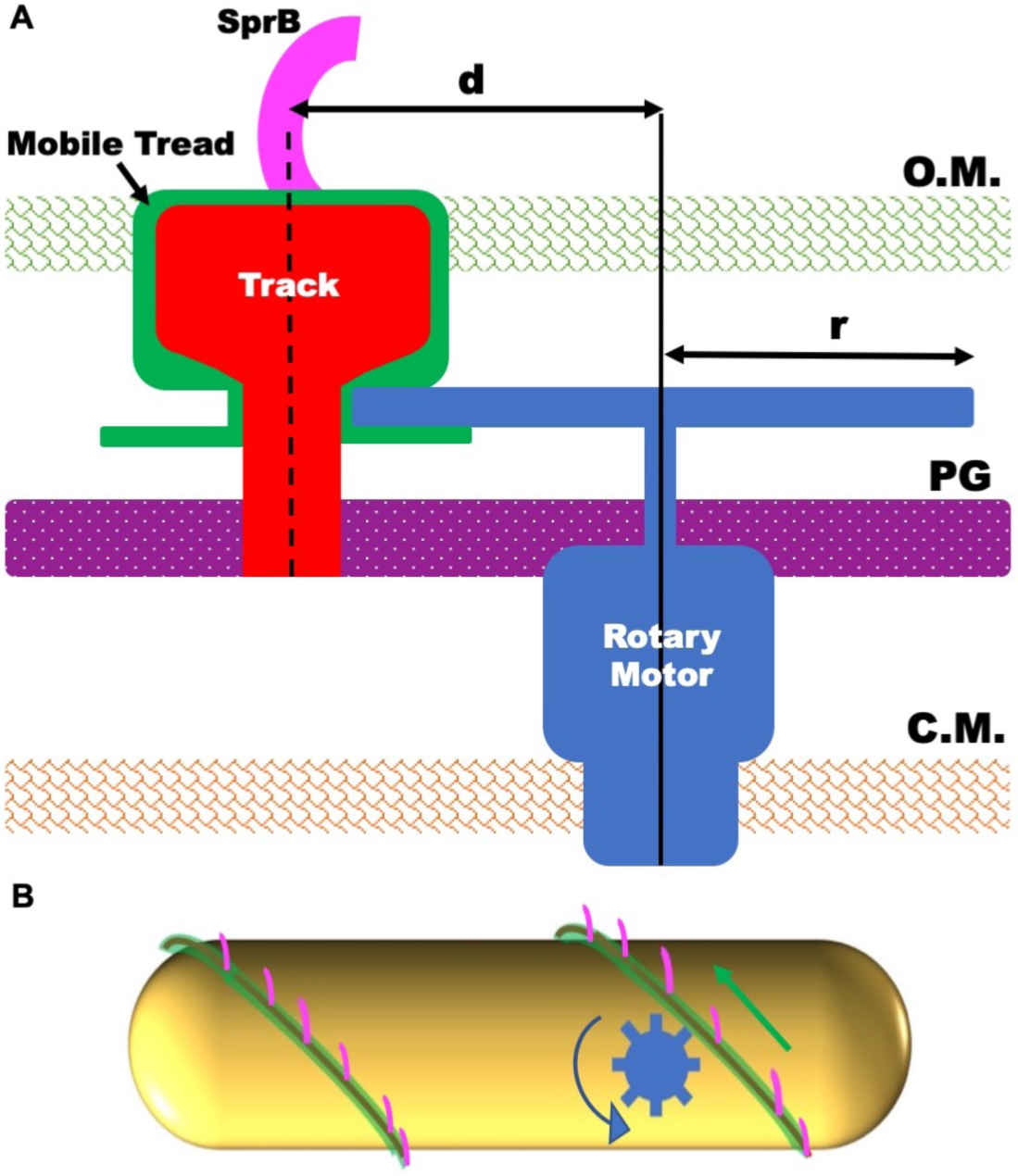
A model of the gliding machinery. **(A)** A cross-sectional view of a cell with a rotary gliding motor (blue), a mobile tread (green), a stationary track (red), and an adhesin (magenta). The rotary motor and the track are anchored to the peptidoglycan (PG), and the track is wound spirally around the cell. The rotary motor drives a pinion that engages a mobile tread (rack) that slides along the track. The adhesin, SprB, is attached to the tread and moves with it. The dimension *d* is the distance between the axis of rotation of the motor and the center of the track, and *r* is the radius of the pinion. **(B)** A side view of a cell with a rotary motor powering the motion of a tread carrying SprB. See Movie 5 for an animation of this model.

The rotary motor, although it runs at constant speed^11^, could be similar in architecture to the flagellar rotary motor, with a rotor embedded in the inner cell membrane surrounded by stator elements (the stationary part of a motor) anchored to the peptidoglycan layer and a drive shaft that passes through the peptidoglycan with the aid of a bushing. The pinion would be mounted on the drive shaft but could rotate under or within the outer cell membrane, driving the tread along a track anchored to the peptidoglycan. Only SprB needs to extend beyond the outer cell membrane. An animation of the behavior of our model is shown in Fig. 4B and Movie 5.

## Discussion

It appears that the presence of multiple SprB’s ensures smooth gliding i.e. if several SprB are mounted on a given tread, a cell can glide continuously. An electron microscope image suggests that there are about 40 SprB on a given cell^10^. So, there will be about 20 on one cylindrical face of a cell. A SprB arriving at the back of a cell can peel off the substrate to be replaced by new SprB adsorbed to the substrate at the front of the cell. As a result, the cell can glide smoothly over many body lengths. However, if a single SprB is mounted on a given tread, that SprB will remain fixed and the free end of the cell will lift off the surface and flip over, allowing the cell to continue to glide in the same direction but without any net progress. Gliding cells often display flips, i.e., the lagging pole of a cell gliding in the x-y image plane gets stuck to glass, the leading pole rises up along the z axis and moves rapidly forming an arc such that it moves behind the lagging pole (Fig. S7 and Movie 5). The flips are illustrated in Fig. 1 and Movie S1 of ref. 7 and in Fig. 4 of ref. 24. After a flip, the direction along which a cell glides changes.

At the time of the early work^23^, no gliding motility proteins were known; all that could be surmised was that sites able to adhere to the substrate moved the length of a cell along tracks fixed to the rigid framework of the cell wall. Fluorescent labeling of proteins that enable gliding motility has improved our knowledge of the mechanism. Future experiments will be aimed towards functional characterization of proteins that form this machinery. Several *F. johnsoniae* mutants are deficient in proteins that either form the T9SS or are involved in the stability of the T9SS proteins. In these mutants, SprB is not secreted to the cell-surface and the cells do not glide. Of more interest are mutants that do secrete SprB to the cell surface but, nonetheless, are not able to glide. An example are cells that lack the C-terminal region of GldJ^24^, a protein that is not a part of the T9SS. Recently, it was found when the C-terminal 8-13 amino acid regions of GldJ, are deleted, secretion via the T9SS is fine while motion of SprB on the cell-surface is significantly reduced GldJ localizes helically, is an outer membrane lipoprotein^25^, and might be associated with the tread/track complex. Identification of additional components of the rack and pinion assembly will enhance our understanding of bacterial gliding.

## Materials and Methods

### Strains and plasmids

A GldL-YFP fusion was generated by amplifying eYFP from pHL55^26^ using primers P65 and P67 (for primers, see Supplementary Information) and cloning it into the vector pCP23. *gldL* was amplified using primers P63 and P64 and was cloned into pCP23 with YFP to generate the plasmid pAS6 such that the C-terminus of GldL was fused with the N-terminus of YFP. Strains were grown at 25°C with shaking, as described previously^1^.

### TIRF imaging of GldL in tethered cells

Cells were tethered using a protocol described previously^11^. To image rotating tethered bacteria with GldL-YFP, a microscope (Nikon Eclipse Ti-U; Nikon, Melville, NJ) with an Apo TIRF 60x 1.49 numerical aperture (NA) objective was used in TIRF mode. A 515 nm, 25 mW laser was used as an excitation source. Fluorescent images were recorded with an Andor Ixon Camera. Phase-contrast images were recorded at the same time in the near infrared using a ThorLabs DCC1545M-GL camera. Images were analyzed using the ImageJ plugin Trackmate^27^ and custom MATLAB codes.

### Simultaneous TIRF imaging of SprB and GldL

SprB was labeled with Cy3 as follows: 2 µL of 1:10 diluted purified anti-SpB antibody^1^ and 2µL of Cy3 conjugated Goat Anti Rabbit IgG Polyclonal Antibody (VWR R611-104-122) were added to 40 µL of 0.4 O.D. bacterial culture and incubated for 10 min at 25°C. After incubation, the preparation was centrifuged at 12,000 × g for 5 min. The supernatant fraction was discarded and the pellet was re-suspended in 40 µL of motility medium (MM; per liter: 1.1 g Casitone, 0.55 g Yeast extract, 1.1 mM Tris (pH 7.5)). A tunnel slide was prepared, 40 µL of 10 mg/mL BSA was added to the tunnel and allowed to stand for 1 min, after which the Cy3-labeled cells were flowed into one end of the tunnel, and a Kimwipe was used to wick the excess fluid from the other end. This preparation was allowed to stand for 10 min, and then was washed twice in the tunnel slide with 40 µL of MM.

To image YFP and Cy3 signals simultaneously, the TIRF system described above was used with a Photometrics DV2 attachment with yellow and red filters (Chroma ET 535/30 and ET 630/75, respectively) to image YFP on one side of the screen and Cy3 on the other. Using a custom MATLAB code, Gaussians with the position along the x-y axis and intensity along the z axis were fit to the Cy3 and YFP spots in each image frame. The position of the peak of each Gaussian fit was recorded as the average position of each spot.

### Addition of CCCP and A22

Wild-type *F. johnsoniae* cells were sheared and tethered to a glass coverslip and attached to a flow cell, and rotation speed was measured as described previously^11^. A CCCP stock was prepared (10 µM in MM and added at the rate of 50 µL/min using a syringe pump, Harvard Apparatus 22). To remove the drug, MM was pumped into the flow cell.

A22 (50 µg/mL), Millipore Sigma, catalogue # 475951, was added to cells gliding on glass in a tunnel slide, as described previously. Cells were imaged using a phase-contrast microscope with a Nikon Plan 40x BM NA 0.65 objective (Nikon Melville, NY) and a Thorlabs DCC1545M-GL (Thorlabs, Newton, NJ) camera. Images were analyzed using a custom MATLAB code.

## Movie Legends

**Movie 1.** A phase-contrast image stack where GldL-YFP strain displays wild-type levels of rotation. Similar to sheared and tethered wild-type cells, two cells in the field of view display rotation. Other cells attach to glass and beads coated with anti-SprB antibody. Some cells and beads display motion over short distances.

**Movie 2.** A tethered cell with 5 fluorescent GldL spots. The cell appears to rotate around a fixed axis.

**Movie 3.** A phase-contrast image stack showing that addition of CCCP stops rotation of a tethered cell.

**Movie 4.** A Cy3-tagged SprB moving along the length of a cell. For this movie, a filter that allows only Cy3 emission to pass was used.

**Movie 5**. An animation of the model suggesting how a molecular rack and pinion could drive gliding. The pinion is blue, the tread green, and the adhesin SprB is magenta.

**Movie 6.** A cell gliding smoothly over a glass surface.

**Movie 7.** A gliding cell displaying flips.

## Author Contributions

AS and HCB designed the experiments and wrote the paper. AS performed the experiments.

## Data and materials availability

All data needed to evaluate the conclusions in the paper are present in the paper and/or the Supplementary Materials. Additional data related to this paper may be requested from the authors.

## Supplementary Information

### Supplementary figure legends

**Figure S1.**
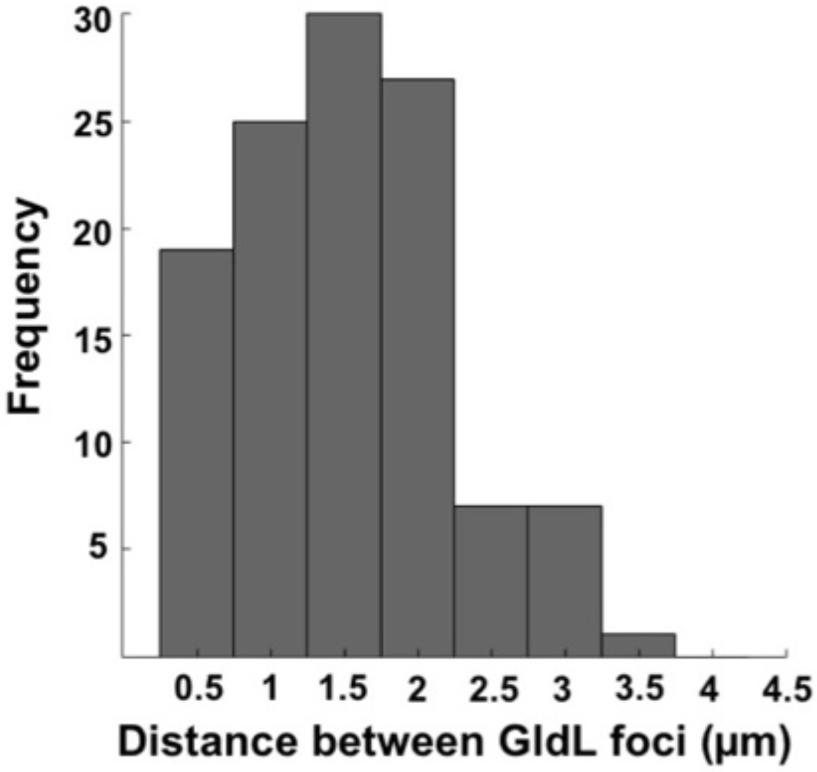
A frequency distribution of nearest-neighbor distances between 42 GldL foci found on 10 cells.

**Figure S2.**
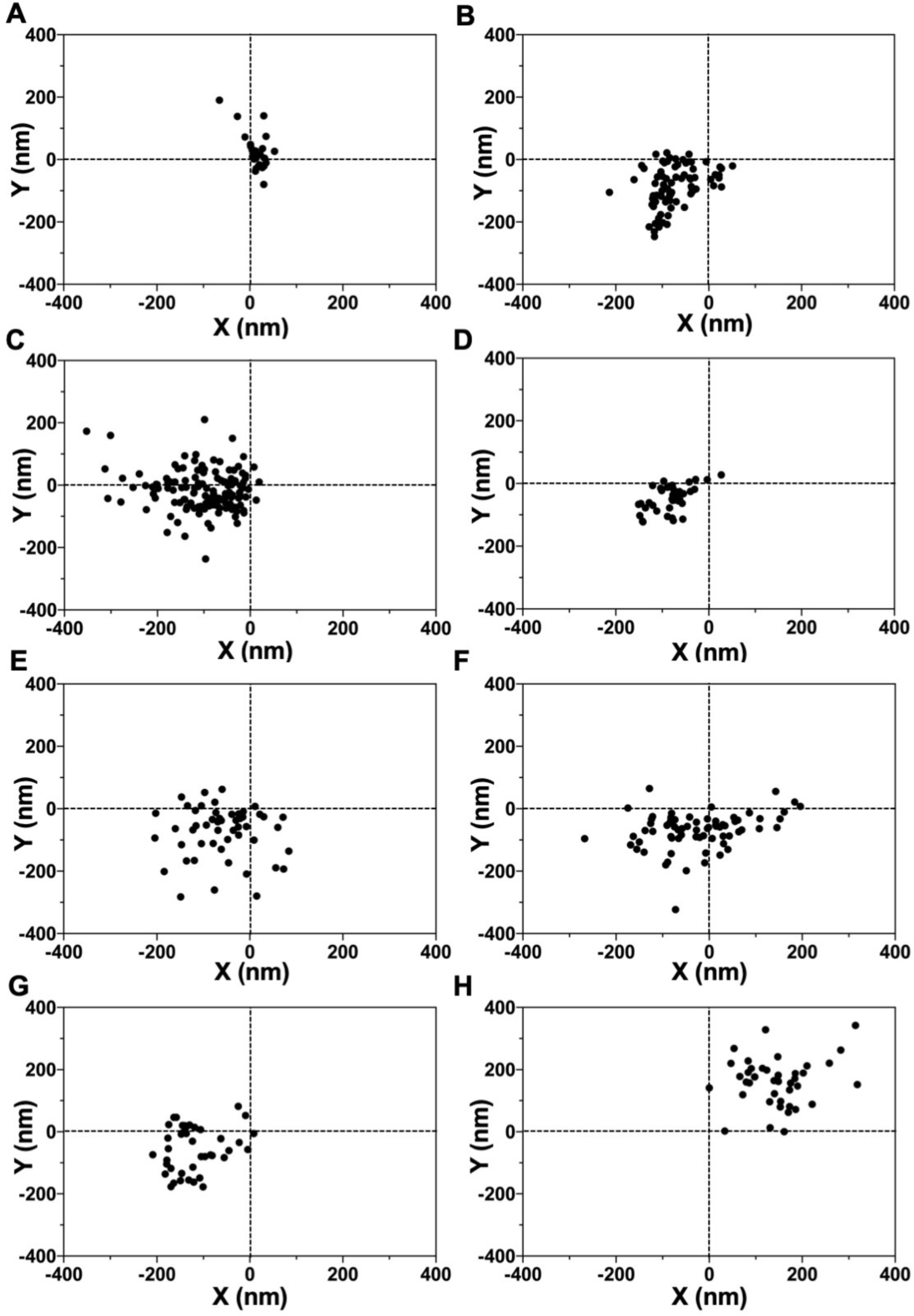
Position of the nearest neighbor GldL foci from the axis of rotation of 8 tethered cells shown for the individual cells. The axis of rotation is normalized to be at the origin. This data are condensed for all 8 cells in Fig. 2A.

**Figure S3.**
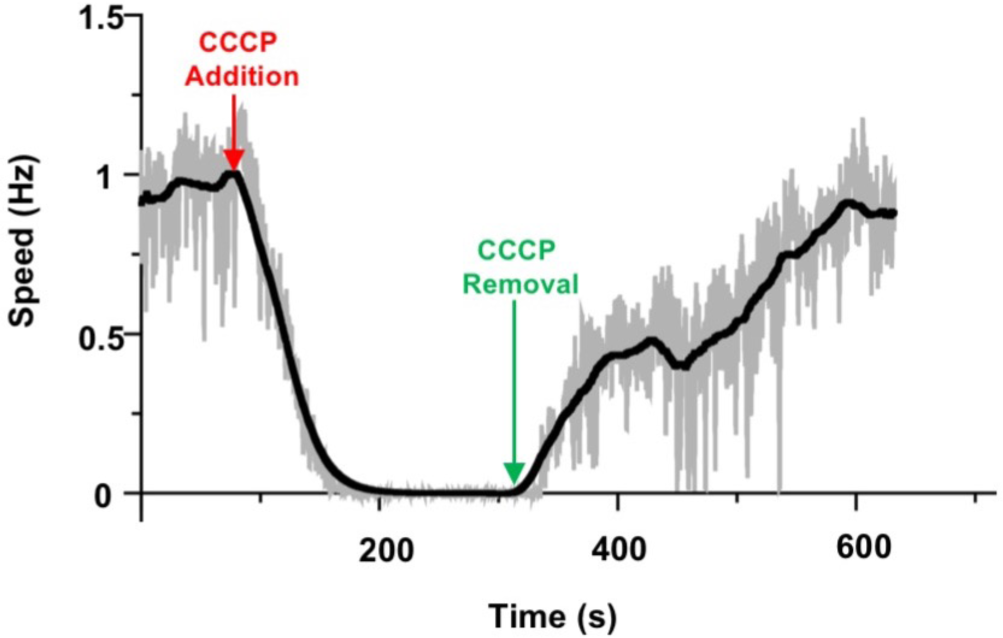
Rotation speed of a cell plotted as a function of time. Addition of CCCP stopped the rotation. Removal of CCCP restored the rotation.

**Figure S4.**
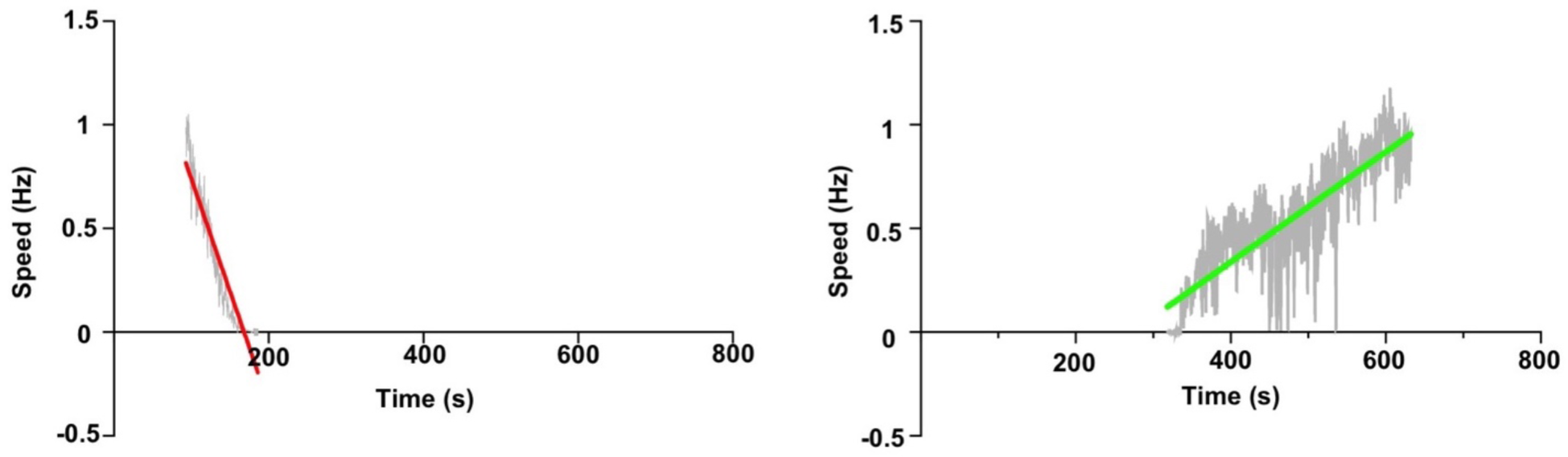
The rate for the ‘drop’ in rotation speed after addition of CCCP (red linear regression fit) is 4 times faster than the rate of increase following removal of CCCP (green fit).

**Figure S5.**
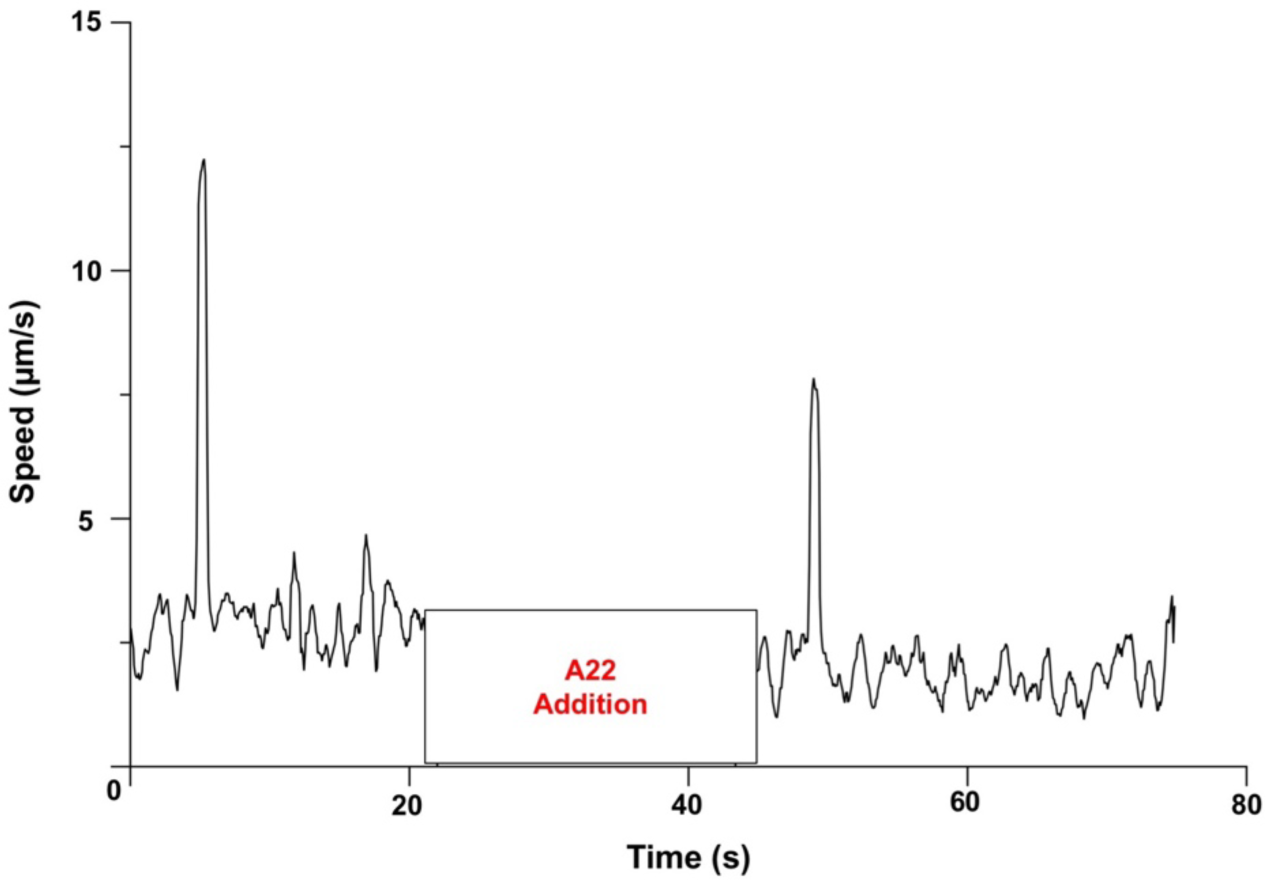
Speed of a smooth gliding cell measured by tracking the center of mass does not change after the addition of A22. The sharp peaks depict flips.

**Figure S6.**
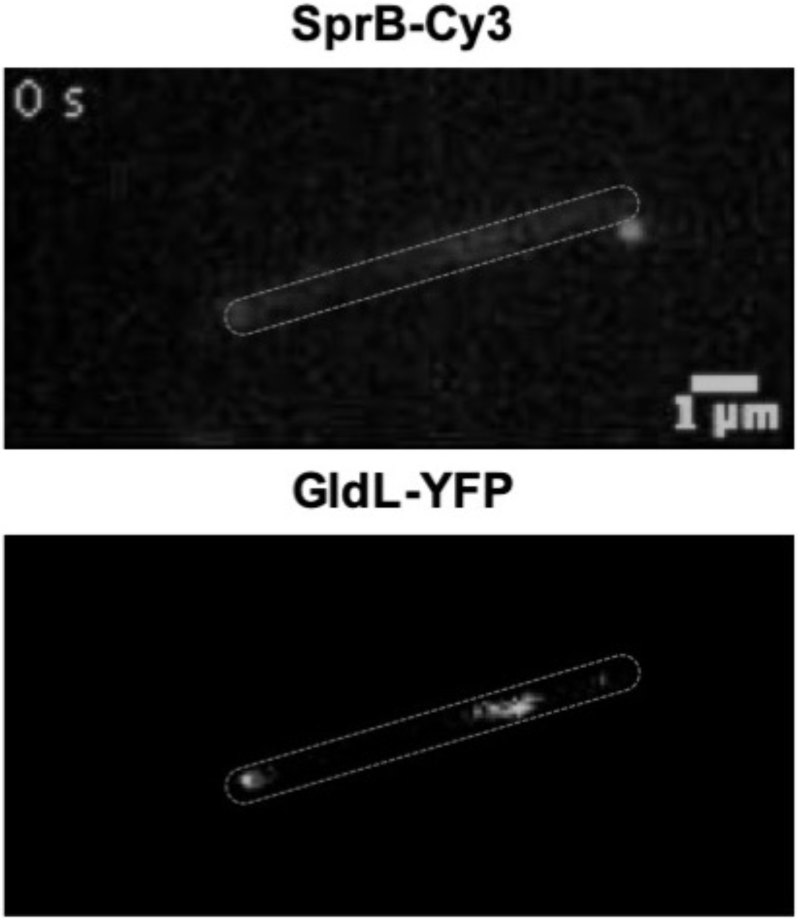
Images of one cell with Cy3 labeled SprB (A) and GldL-YFP foci (B). Cell boundary is marked with a dotted line. Panel A is an image of a cell where a filter that allows only Cy3 emission to pass was used. Panel B is an image of the same cell where a filter that allows only YFP emission to pass was used.

**Figure S7.**
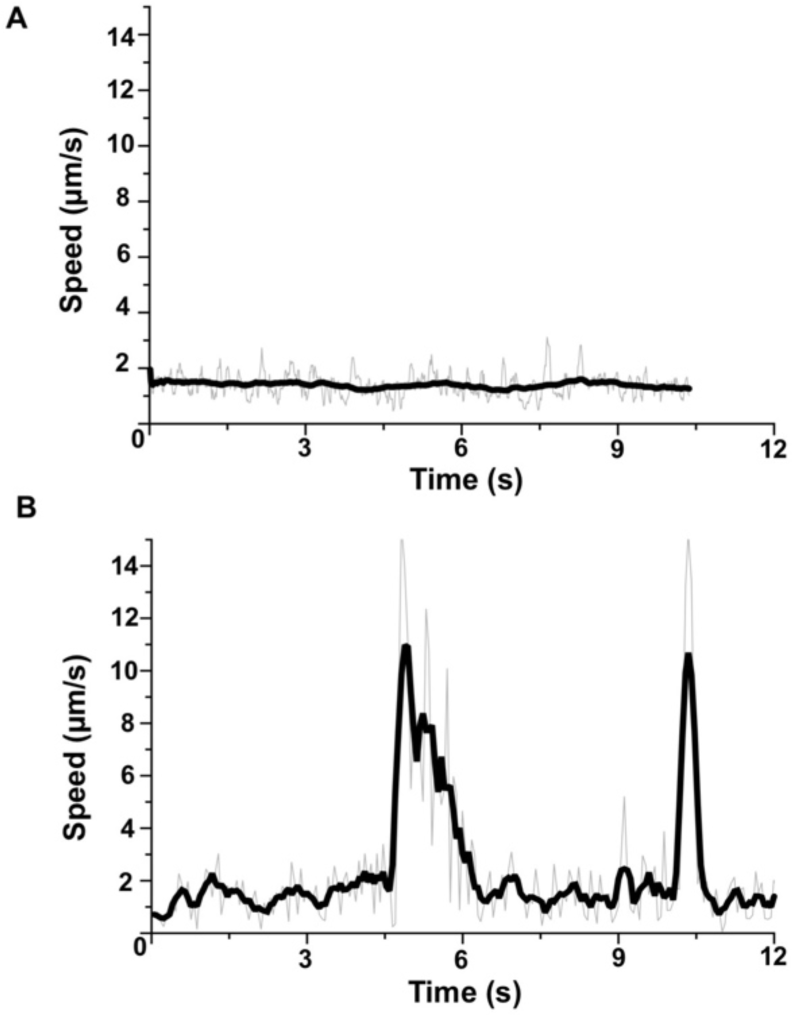
Tracking the flips displayed by gliding cells. **(A)** Speed of a smooth gliding cell measured by tracking the center of mass of a gliding cell when plotted as a function of time does not change (Movie 6). **(B)** The speed of cell that displays two flips when plotted as a function of time show two sharp peaks (Movie 7).

## Primers used in the study

### P63

5’ GCTAG<u>GGTACC</u>GGAACTCAAGTAACAGGAGGCG 3’ forward primer upstream of gldL, KpnI site underlined.

### P64

5’ GCTAG<u>GGATCC</u>TCCTTTGTTACTCATTGCAGAAAG 3’ reverse primer in gldL, BamHI site underlined.

### P65

5’GCTAG<u>GGATCC</u>**GGAGGTGGA**ATGGTGAGCAAGGGCGAG 3’ Binds YFP start site in pHL55. BamHI site underlined, 3xG linker in bold.

### P67

5’ GCTAG<u>TCTAGA</u>**TTA**CTTGTACAGCTCGTCCATGCCGAGAGTGATCCCGGCGGC 3’

Binds end region of yfp. Stop codon in bold. XbaI site underlined. *gldL-YFP* in the plasmid pCP23 was confirmed by sequencing.

### Computer Scripts

Custom MATLAB codes used to analyze the data for this article are freely available on GitHub https://github.com/Abhishek935/Molecular-Rack-and-Pinion.

